# Cas9-based enrichment for targeted long-read metabarcoding

**DOI:** 10.1101/2024.12.02.626365

**Authors:** Lucia Nikolaeva-Reynolds, Christopher Cammies, Rosemary Crichton, Thomas E. Gorochowski

## Abstract

Metabarcoding is a valuable tool for characterising the communities that underpin the functioning of ecosystems. However, current methods often rely on PCR amplification for enrichment of marker genes. PCR can introduce significant biases that affect quantification and is typically restricted to one target loci at a time, limiting the diversity that can be captured in a single reaction. Here, we address these issues by using Cas9 to enrich marker genes for long-read nanopore sequencing directly from a DNA sample, removing the need for PCR. We show that this approach can effectively isolate a 4.5 kb region covering partial 18S and 28S rRNA genes and the ITS region in a mixed nematode community, and further adapt our approach for characterising a diverse microbial community. We demonstrate the ability for Cas9-based enrichment to support multiplexed targeting of several different DNA regions simultaneously, enabling optimal marker gene selection for different clades of interest within a sample. We also find a strong correlation between input DNA concentrations and output read proportions for mixed-species samples, demonstrating the ability for quantification of relative species abundance. This study lays a foundation for targeted long-read sequencing to more fully capture the diversity of organisms present in complex environments.

## INTRODUCTION

The characterization of micro-organisms and meso-faunal communities is a powerful tool in environmental, agricultural, medical, and biotechnological settings due to the essential roles these organisms play across diverse biological systems [1, 2]. For example, nematodes are excellent bioindicators of soil health and quality, and can be used to accurately predict primary decomposition pathways [3, 4]. The main challenge to unlocking the use of this community knowledge is accurately identifying the often large numbers of species that are present and quantifying their relative abundance. Unfortunately, traditional methods like morphological identification are time-consuming and requires a high level of taxonomic expertise. This puts it outside the reach of most studies and limits the depth of analysis that can be performed [5].

Metabarcoding is a powerful alternative to traditional methods of identification that typically require visual observations and/or physical separation of individual organisms prior to further analysis (e.g., gram staining of bacteria). Metabarcoding typically uses polymerase chain reaction (PCR) and high throughput sequencing to simultaneously amplify and sequence molecular barcodes, such as the commonly used cytochrome c oxidase subunit I (COI) [6–8], 16S or 18S ribosomal (r)RNA genes [9, 10], and internal transcribed spacer (ITS) region [11, 12], from a metagenomic sample [13, 14]. This allows for the identification of many species simultaneously from mixed species samples, like those taken from water or soil [8–10, 15]. Metabarcoding has been valuable in answering ecological questions such as ecosystem biomonitoring [7, 8, 16], revealing dietary profiles using fecal DNA [17, 18], reconstructing food webs [19] and ancient community dynamics from relic DNA [20].

Whilst a useful tool, PCR-based metabarcoding does have limitations. Firstly, the nucleotide composition of template DNA can influence PCR efficiency. For example, homopolymers, GC-rich regions, and inverted repeats can be problematic for amplification [21–24]. Also, stochastic variation early on in the PCR process can lead to significant bias in the final template-to-product ratio [21, 25], causing skewed estimates of relative abundance of different species [21]. Furthermore, current PCR-based barcoding is often limited by the taxonomic resolution achievable through the use of a single barcode region (typically 200–600 bp long), which captures only partial variation of interspecific nucleotides along these genes [26, 27].

Metabarcoding studies that aim to capture phylogenetically diverse communities may use a ‘universal barcode’ such as the COI gene [6]. However, the true ‘universality’ of universal barcodes is contested [28], where amplification success rates are low for certain taxonomic groups, skewing diversity estimates [27, 29, 30]. Due to these issues, metabarcoding studies that want to characterise complex communities often face a trade-off between diversity and resolution. To address this, multiple PCR assays can be performed to target different taxa within the sample, using optimised primer pairs for each target [12, 16]. However, this is time-consuming and hampers comparisons of abundance between taxa.

Third-generation sequencing platforms, from Oxford Nanopore Technologies (ONT), PacBio, Quantapore and Stratos, provide an alternative to amplification-based DNA sequencing. These technologies can sequence native DNA and RNA, eliminating the need for PCR. The long-read capabilities of these third-generation platforms also allow for longer barcode regions to be sequenced, capturing a greater proportion of interspecific nucleotide variation. This increases taxonomic resolution and allows for species, or even strain, identification [31–33]. Nanopore sequencing has been applied to microbial, nematode, and vertebrate diversity studies [34–38]. There are two studies to date that use nanopore sequencing for nematode identification, both of which use individual organisms as starting material [37, 39]. Nanopore sequencing has not yet been used to identify nematode species from metagenomic samples, as has been done for microbes [38]. Although not a requisite for nanopore sequencing, all of these long-read based studies use PCR as a mode of enrichment of target genes and so still face issues in quantifying diverse communities [36, 38].

A novel approach to enrichment of specific nucleotide sequences employs the use of clustered regularly interspaced short palindromic repeats (CRISPR) systems. CRISPR naturally occurs as a defence system in bacteria and archaea against foreign viruses and plasmids using nucleases such as Cas9 to target and cleave specific DNA sequences [40, 41]. The ability to easily target DNA cleavage via CRISPR (cr)RNAs has led to CRISPR becoming widely adopted as a flexible tool for genome editing [42, 43]. CRISPR-based technology has also been used to isolate specific regions of native DNA for sequencing, removing the need for enrichment via PCR. [44]. This approach has been shown to be compatible with nanopore sequencing, and has been used to isolate the human breast cancer gene BRCA1, to characterise genetic variants where the region of interest was excised and then isolated using pulsed-field gel electrophoresis (PFGE) [45].

More recently, a targeted nanopore sequencing approach using Cas9 was proposed [46]. This method relies on cleavage of native target DNA by the Cas9 enzyme [47] and attachment of sequencing adapters to the cleaved target DNA **(Figure 1A)**. To date, Cas9-based enrichment has been used to increase sequencing coverage at specific regions of interest to assess single nucleotide polymorphisms [48], structural variations [48], repetitive regions [49, 50] and epimutations [50]. This method of enrichment also enables multiple DNA targets to be enriched simultaneously by multiplexing (i.e., combining) crRNA probes that guide cleavage to multiple target regions in a single library preparation. This contrasts with PCR amplification as PCR requires specific annealing temperatures for each PCR primer, making it unfeasible to include multiple primers that target different regions. While this type of enrichment has been shown to be a promising approach for understanding specific genetic features, it has yet to be applied to metagenomic studies.

**Figure 1:**
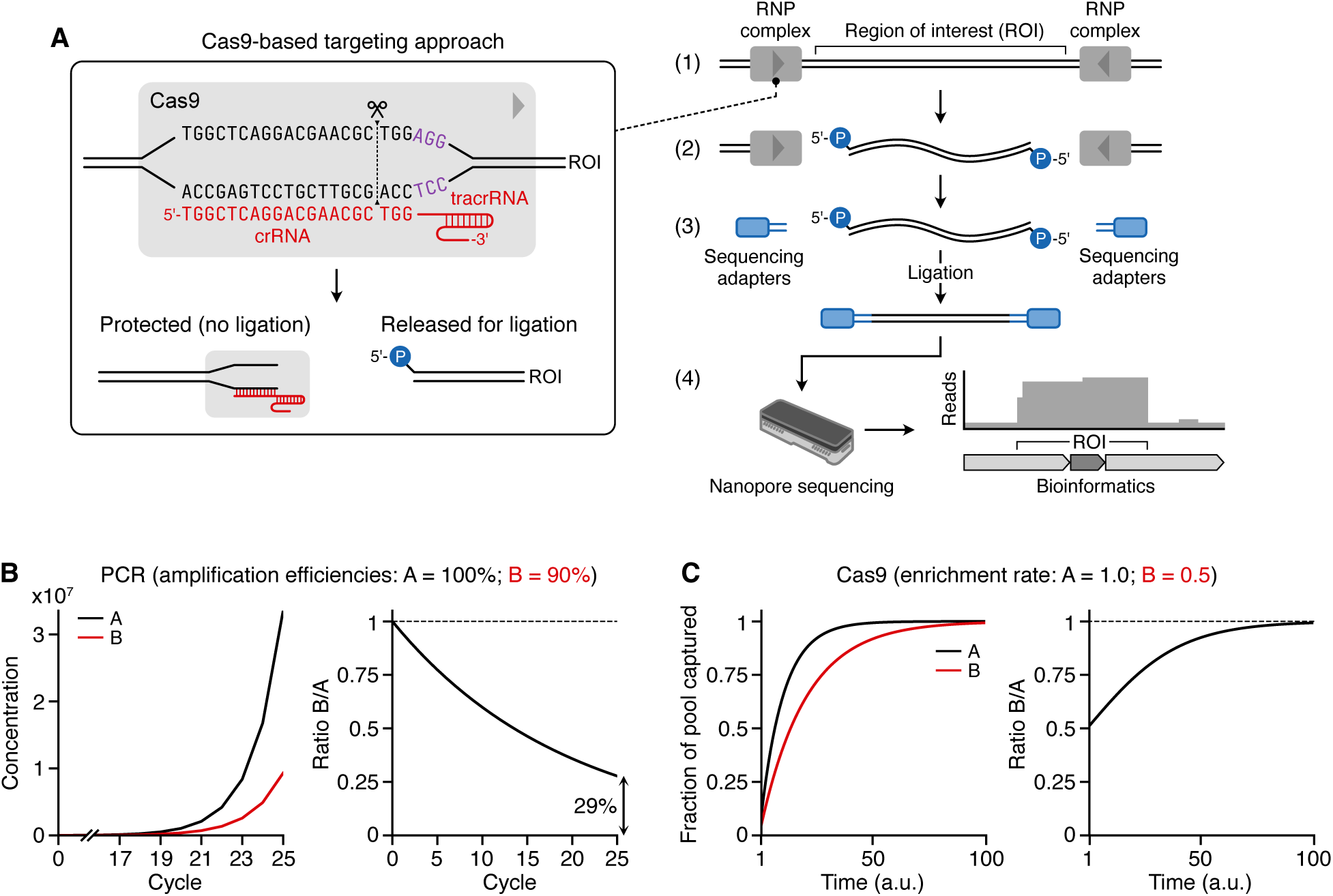
Overview of the Cas9-based targeted sequencing approach. (**A**) Major steps in the Cas9-based enrichment: 1. Targetting of Cas9 using crRNAs that bind to flanking points of a region of interest (ROI); 2. Cleavage of sample DNA and protection of fragment ends that should not be enriched; 3. Ligation of sequencing adapters to unprotected ends of DNA fragments from the ROI; 4. Nanopore sequencing and bioinformatics to recover enriched data for the ROI. (**B**) PCR amplification model. Left plot shows expected PCR amplification of two different products ‘A’ (black) and ‘B’ (red) that start at identical concentrations of 1 molecule/sample each using primers that are 100% and 90% efficient, respectively. Right plot shows the ratio B/A and large deviation in the expected 1:1 ratio (doted line) after 25 cycles of PCR. (**C**) Cas9-based enrichment model where Cas9-gRNA complexes are assumed in excess. Left plot shows expected enrichment (as a fraction) of two different products ‘A’ (black) and ‘B’ (red) that have an equal concentration in the sample, but where the rate of cleavage of the Cas9-gRNA complex (i.e., enrichment) in B is half that of A (rates of 0.5 and 1.0, respectively). Right plot shows the ratio B/A and the strong trend towards the expected ratio of 1 (dotted line) as the length of the reaction increases.

In this paper, we address this gap and apply targeted nanopore sequencing in a metabarcoding context to detect and quantify species in singular and mixed species samples. We demonstrate that our method of amplification-free metabarcoding allows us to target multiple barcode regions across taxonomic groups to more fully characterize diverse communities in a single reaction. This removes the need to carry out multiple PCR reactions to fully uncover diversity, saving considerable time and resources. Furthermore, the lack of PCR helps to reduce potential biases surrounding differences in amplification efficiency between taxonomic groups, allowing for improved estimates of relative abundance. We see this method as a further tool for biologists to better understand the true diversity of life in complex environments, exploiting the ability for amplification-free, long-read metabarcoding to enhance the breadth and depth of community profiling.

## RESULTS

### Cas9-based enrichment for metabarcoding

Targeted nanopore sequencing using Cas9-based enrichment works by cleaving target DNA sequences at user defined points and attaching nanopore sequencing adapters to one of the cleaved ends (**Figure 1A**) [46]. Prior to cleavage, the ends of extracted genomic DNA are dephosphorylated to avoid ligation of nanopore sequencing adapters to non-target DNA that may have been fragmented during extraction. Then, custom 20-mer CRISPR (cr)RNA sequences (“probes”) are combined with catalytic trans-activating CRISPR (tracr)RNA, forming a guide (g)RNA targeting mechanism for the Cas9 nuclease, which can then bind and cleave at sequences identical or highly similar to the crRNA. The 20-mer target site (the protospacer) must be adjacent to an NGG-sequence protospacer-adjacent motif (PAM) for the Cas9 to function efficiently. The Cas9 cleaves the target sequence 3 bp upstream of the PAM. These cuts expose a 5’ phosphate group, onto which nanopore sequencing adapters can be ligated, however, the Cas9-gRNA complex also block ligation to the end of one of the fragments (the non-targeted region). Therefore, the resulting library contains DNA molecules where sequencing adapters are mostly attached to target DNA. This enriched library results in a greater ratio of target to non-target DNA being sequenced than if no enrichment took place.

A potential benefit of using Cas-based enrichment rather than PCR amplification for targeted metabarcoding is that small differences in the amplification efficiency of primers used during PCR to target barcode regions across species can lead to large deviations in the recovered abundances. To demonstrate this, we developed idealised mathematical models of the two enrichment methods. In both, we assumed two different target species (‘A’ and ‘B’) with identical concentrations, but where amplification or enrichment efficiency differed between the species. For PCR amplification (**Figure 1B**), if the efficiency of amplification of B is only 10% lower than for A, then after 25 cycles, the original ratio of B/A = 1 will have fallen to 0.29. In contrast, Cas9-based enrichment (**Figure 1C**) is less affected by differences in the efficiency of cleavage for crRNAs in the target species. If the Cas9-gRNA complex is in excess for any enrichment reaction, and assuming the reaction is run for a sufficiently long time, then even when there are large differences in cleavage efficiency (e.g., cleavage rate of B is half that of A), correct ratios of the underlying species can still be recovered (**Figure 1C**).

To evaluate the potential of Cas9-based enrichment for targeted metabarcoding, we tested its ability to enrich typical barcode regions in the nematode *Caenorhabditis elegans*. We designed crRNA probes to target a ∼4.5kb region within the rDNA tandem repeat. The target region included partial 18S rDNA (v6–v8 regions), 28S rDNA (D1–D10 regions), both ITS regions and the complete 5.8S rDNA gene. We applied the enrichment strategy to extracted *C. elegans* genomic (g)DNA (**Figure 2**; **Methods**). To measure enrichment, we calculated an enrichment score, *E*, representing the fold-increase in reads covering the target sequence compared to the expected coverage if reads were evenly distributed across the genome (i.e., without enrichment). This can be calculated using,

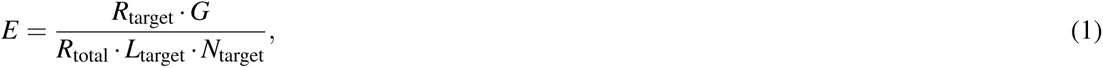

**Figure 2:**
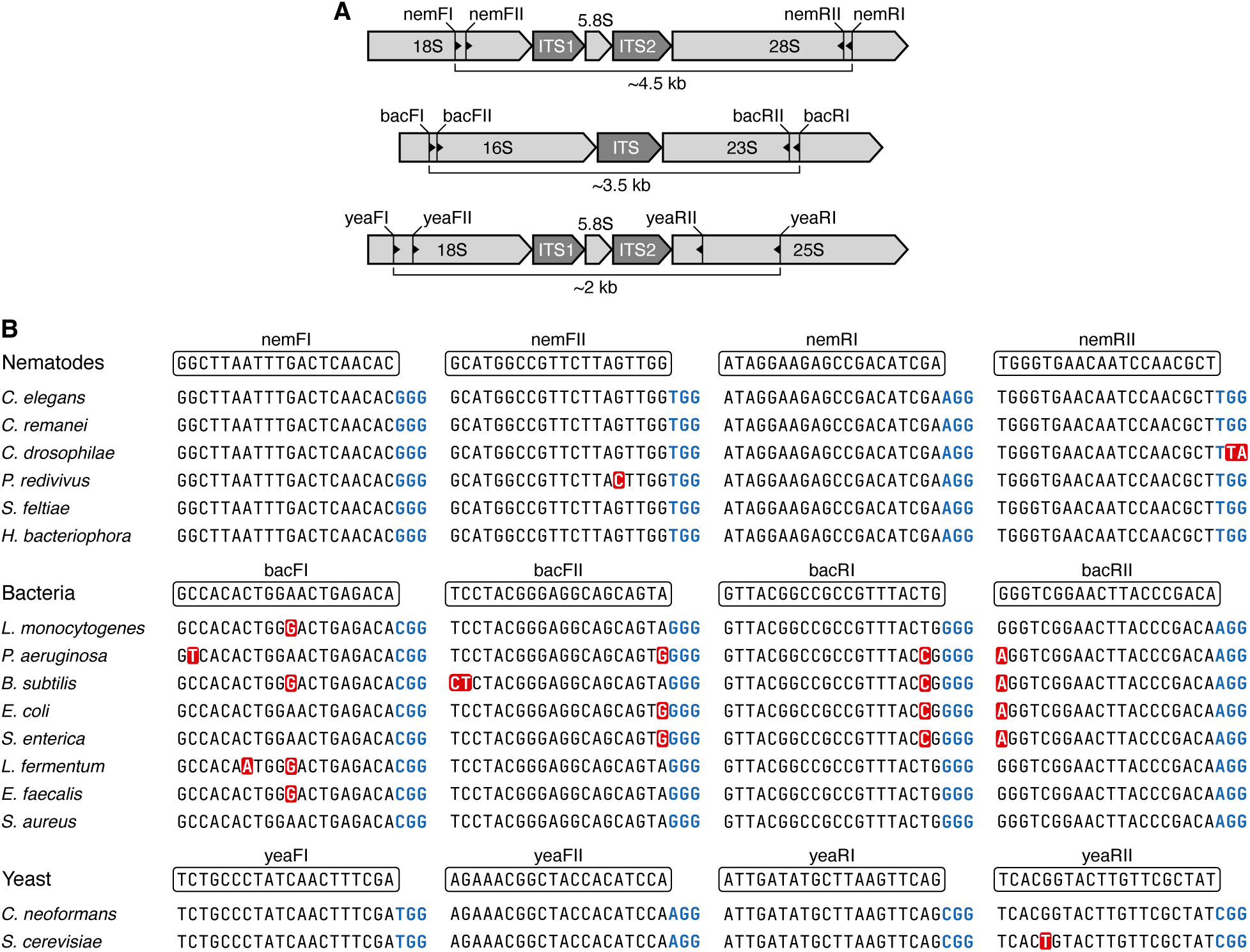
Design of crRNA probes targeting rRNA encoding genomic regions. (**A**) crRNA probe locations within the rRNA encoding genomic regions for nematodes (top), bacteria (middle), and yeast (bottom). Direction of the crRNA shown by small black triangle. (**B**) Alignment of crRNA sequences to corresponding genomes. Sequences for species without documented genomes use 18S (for forward crRNAs) and 28S (for reverse crRNAs) sequences for this alignment (**Data S1**) as the crRNAs sit within these barcodes. Mismatches to crRNA (boxed sequence) highlighted in red. Note that the Cas9 ‘NGG’ protospacer adjacent motif (PAM) sequence (bold, blue) is not part of the crRNA sequence.

where *R*_target_ is number of on-target reads, *R*_total_ is total number of reads, *G* is genome size, *L*_target_ is length of the target region, and *N*_target_ is number of copies of the target region (see **Table 1** for genome sizes and copy numbers).

**Table 1:** Organisms used in the mock communities ^a^.

| Species | Type | Genome size (Mbp) | gDNA com- position by mass (%) | rDNA copies per genome | Abundance % based on 16S & 18S<sup>b</sup> |
| --- | --- | --- | --- | --- | --- |
| *Caenorhabditis elegans* | Nematode | 100.2 | Variable<sup>c</sup> | 55<sup>d</sup> | — |
| *Caenorhabditis remanei* | Nematode | 124.8 | Variable<sup>c</sup> | 183<sup>d</sup> | — |
| *Caenorhabditis drosophilae* | Nematode | 51.3 | Variable<sup>c</sup> | — | — |
| *Heterorhabditis bacteriophora* | Nematode | 77 | Variable<sup>c</sup> | — | — |
| *Steinernema feltiae* | Nematode | 91 | Variable<sup>c</sup> | — | — |
| *Panagrellus redivivus* | Nematode | 65 | Variable<sup>c</sup> | — | — |
| *Listeria monocytogenes* | Bacteria | 2.99 | 12 | 6 | 12.4 |
| *Pseudomonas aeruginosa* | Bacteria | 6.8 | 12 | 4 | 3.6 |
| *Bacillus subtilis* | Bacteria | 4.0 | 12 | 10 | 15.3 |
| *Escherichia coli* | Bacteria | 4.9 | 12 | 7 | 8.9 |
| *Salmonella enterica* | Bacteria | 4.8 | 12 | 7 | 9.1 |
| *Lactobacillus fermentum* | Bacteria | 1.9 | 12 | 5 | 16.1 |
| *Enterococcus faecalis* | Bacteria | 2.8 | 12 | 4 | 8.7 |
| *Staphylococcus aureus* | Bacteria | 2.7 | 12 | 6 | 13.6 |
| *Saccharomyces cerevisiae* | Yeast | 12.1 | 2 | 109 | 9.3 |
| *Cryptococcus neoformans* | Yeast | 18.9 | 2 | 60 | 3.3 |

a. Separate communities studied are grouped in the table and unknown quantities are denoted with a dash.
b. Calculated based on rDNA copy number and genome size given by Zymobiomics.
c. For details see **Data S2**.
d. rDNA copy numbers taken from [75] for *C. elegans* and [76] for *C. remanei*.

We obtained between 0.46 M and 1.64 M reads from the sequencing experiments, with the number of reads increasing with the starting DNA mass (**Data S2**). A high level of enrichment of the target region was observed for all samples. A mean of 83.3% of reads were on target, with an average enrichment of *E* = 337-fold that was consistently high across all replicates (*E* = 348, 344, 318-fold for replicate samples 1a–c, respectively) **(Figure 3)**. Read coverage of the *C. elegans* genome increased sharply at the beginning of the target region and decreased sharply at the end of the target region, points where the ‘forward’ and ‘reverse’ crRNA probes guided cleavage, respectively **(Figure 3)**. Coverage for forward and reverse strands were approximately equal for all replicates, demonstrating equal activity of forward and reverse probes (**Figure 3**). A distinct peak of ∼4.5 kb was observed in the raw read lengths in all three replicates, matching the anticipated fragment lengths from simulated Cas9 digestion (**Figure 4**).

**Figure 3:**
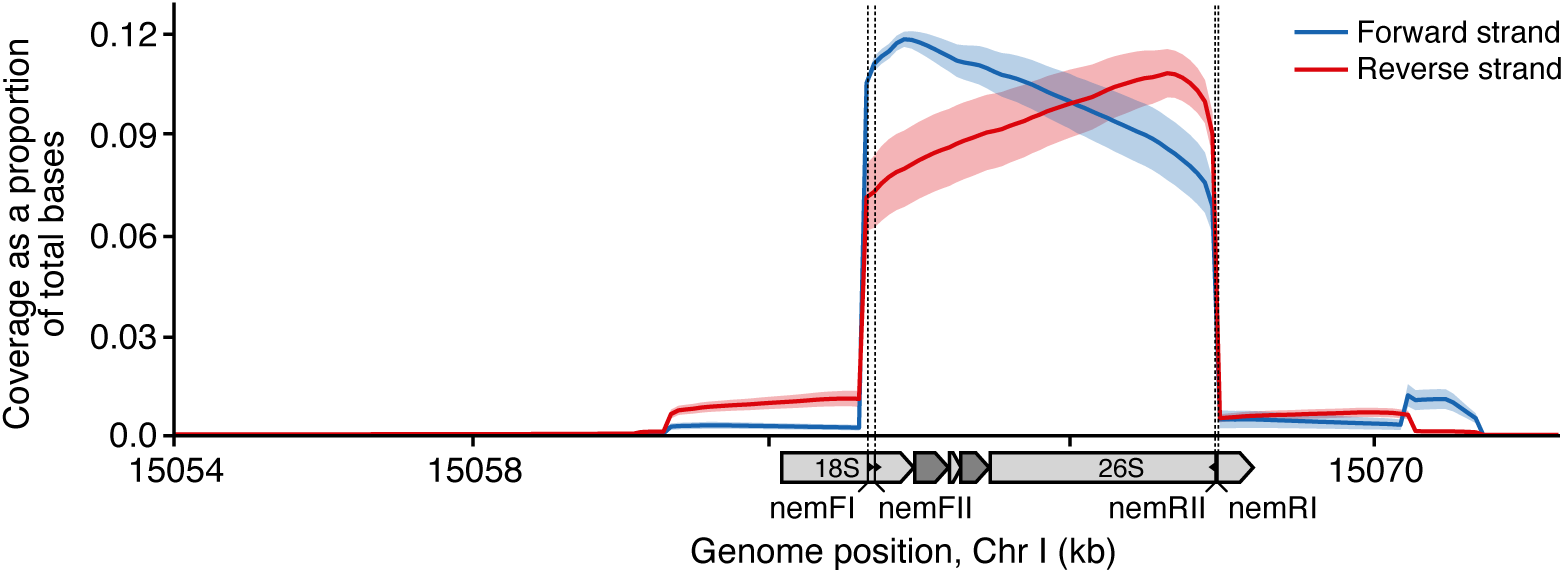
Coverage plots of *C. elegans* depicting enrichment of target region. Coverage for targeted sequencing of a pure *C. elegans* samples given as the proportion of total bases per sequencing run that map to a nucleotide position. Target region covers genomic coordinates chrI:15063336-15067815 in the *C. elegans* genome, marked by the positions of the crRNAs: nemFI, nemFII, nemRI, and nemRII. Solid lines represent the mean coverage across three replicates and shaded area denotes the absolute deviation from the mean.

**Figure 4:**
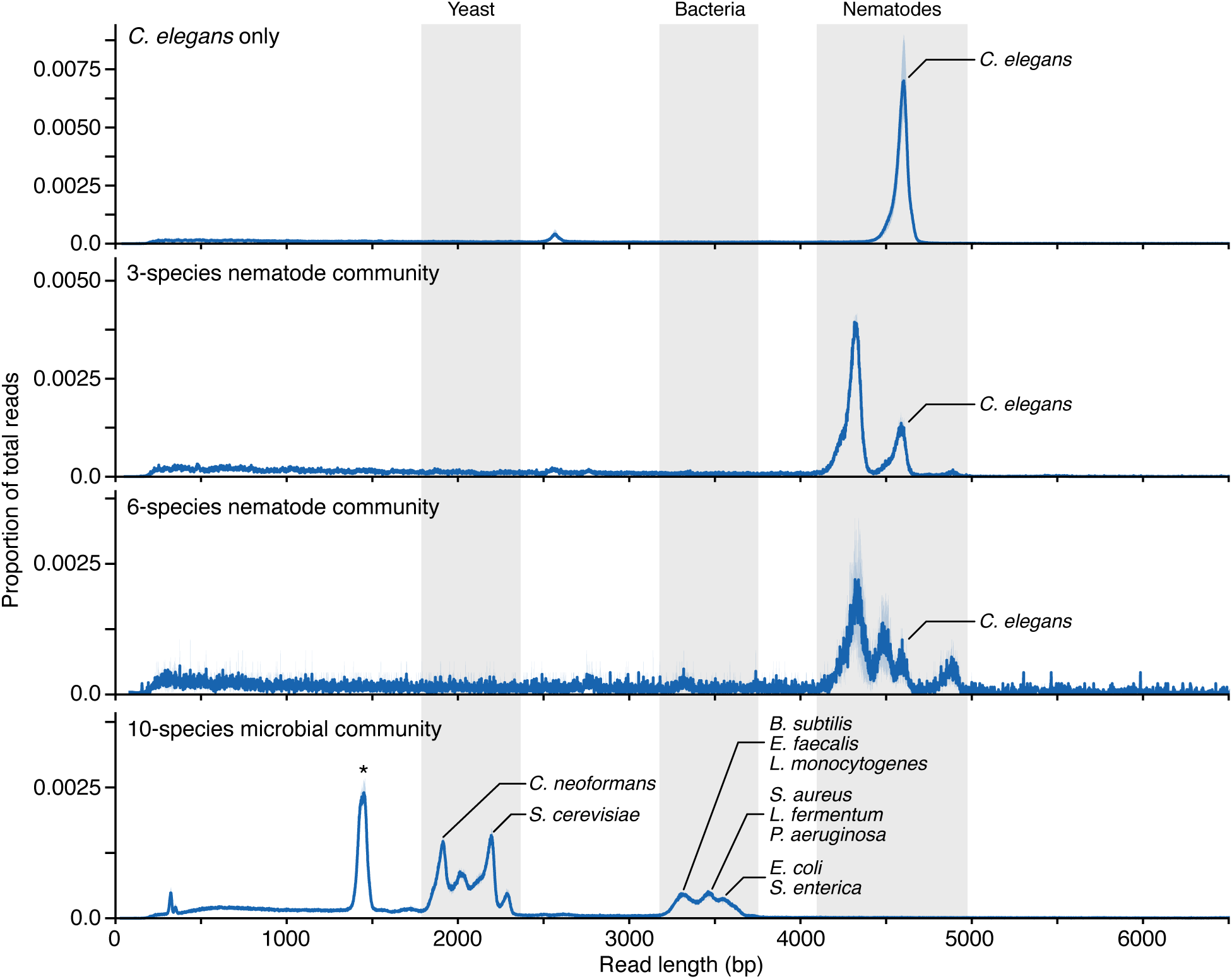
Read length distributions for the four communities assessed. Blue solid lines show the average of the three replicates and shaded blue regions denotes the absolute deviation from the mean. Grey shaded regions across samples correspond to the expected target lengths of the rRNA regions in bacteria, yeast and nematodes. Peaks for specific species have been annotated. The peak denoted with a star marks unexpected 1.5 kb reads which are bacterial reads cleaved by off-target activity of yeast crRNAs.

### Targeted nanopore sequencing of mock nematode communities

Having demonstrated that Cas9-based enrichment is effective for targeted sequencing of a barcode region in *C. elegans*, we next applied the same set of crRNA probes to metagenomic samples to test whether multiple species could be identified using this method. Two sets of custom mock nematode communities were used; one consisting of three species: *C. elegans*, *Heterorhabditis bacteriophora*, and *Steinernema feltiae* (Samples 2a–c), and the other consisting of six species: *C. elegans*, *H. bacteriophora*, *S. feltiae*, *Caenorhabditis remanei*, *Caenorhabditis drosophilae*, and *Panagrellus redivivus* (Samples 3a–c).

Distinct peaks between 4.2 kb and 4.9 kb were observed in the read length distributions for all nematode sequencing runs (**Figure 4**), confirming the expected fragment lengths for the enrichment protocol. As the samples become more diverse in species, peaks become more numerous around the 4.5 kb mark due to interspecific variation in rDNA length, demonstrating the generation of target read lengths for the multiple species (**Figure 4**).

Sequencing data was further analysed using a metagenomics workflow (**Methods**). When using a 98% similarity threshold between nanopore reads and reference sequences, all nematodes were detected to genus level in all replicates for both the three- and six-species communities. Between two to four extra genera were identified in the analysis of each dataset, but falsely positive genera had read abundances of *<*0.08% of the total mapped reads, and most had only a single read classified to the genus. When analysis was done to a species-level, many false positives occurred, and up to 31 species were classified. Total input DNA ranged from 510–2917 ng for the mock nematode communities (**Data S2**), and the number of reads generated from sequencing increased as the amount of input DNA increased.

### Multiplexing crRNA probes to characterise complex microbial communities

To analyse the relative abundance of taxa in a sample using DNA marker abundance, copy numbers of the genes being used are required to account for repeats of the target gene within a genome [51]. As this information is not known for all nematode species used in this study, quantitive analysis could not be performed for the nematode mock communities. To overcome this limitation, we applied our method to the Zymo microbial community DNA standard. This contains ten species of bacteria and yeast in known gDNA proportions with defined marker gene abundance, and available genome sequences **(Table 1)**. We multiplexed two sets of crRNA probes in a single library preparation, one set targeting the rDNA region in bacteria and the other set the rDNA region in yeast. The target regions included commonly used 16S, 18S and ITS barcode regions (**Figure 2**; **Methods**).

Despite long sequencing times, the generation of reads tended to plateau after 12–24 hours, depending on the amount of input DNA, and qualitative and quantitative results were consistent across samples despite differences in input DNA mass and sequencing times (**Data S2**). Distinct peaks were observed for the expected read lengths of each taxon; ∼1.9 kb in *C. neoformans*, ∼2.2 kb in *S. cerevisiae*, and ∼3.3–3.7 kb in bacterial species (interspecific variation of rDNA lengths resulted in multiple peaks in the read length distributions) (**Figure 4**; **Table 2**). A large unexpected peak was also observed around 1.4 kb (**Figure 4**). Further investigation of this peak found that there is a small region in the bacterial genome, 1.4 kb upstream of the bacRII probe, with sequence similarity to the yeaRII probe, albeit with two mismatches. It is therefore likely that the peak is caused by off-target activity of the yeast probe, cleaving the target reads of the bacteria. Nevertheless, 1.4 kb of bacterial rDNA sequence is ample for taxonomic classification, and did not seem to skew results.

**Table 2:**
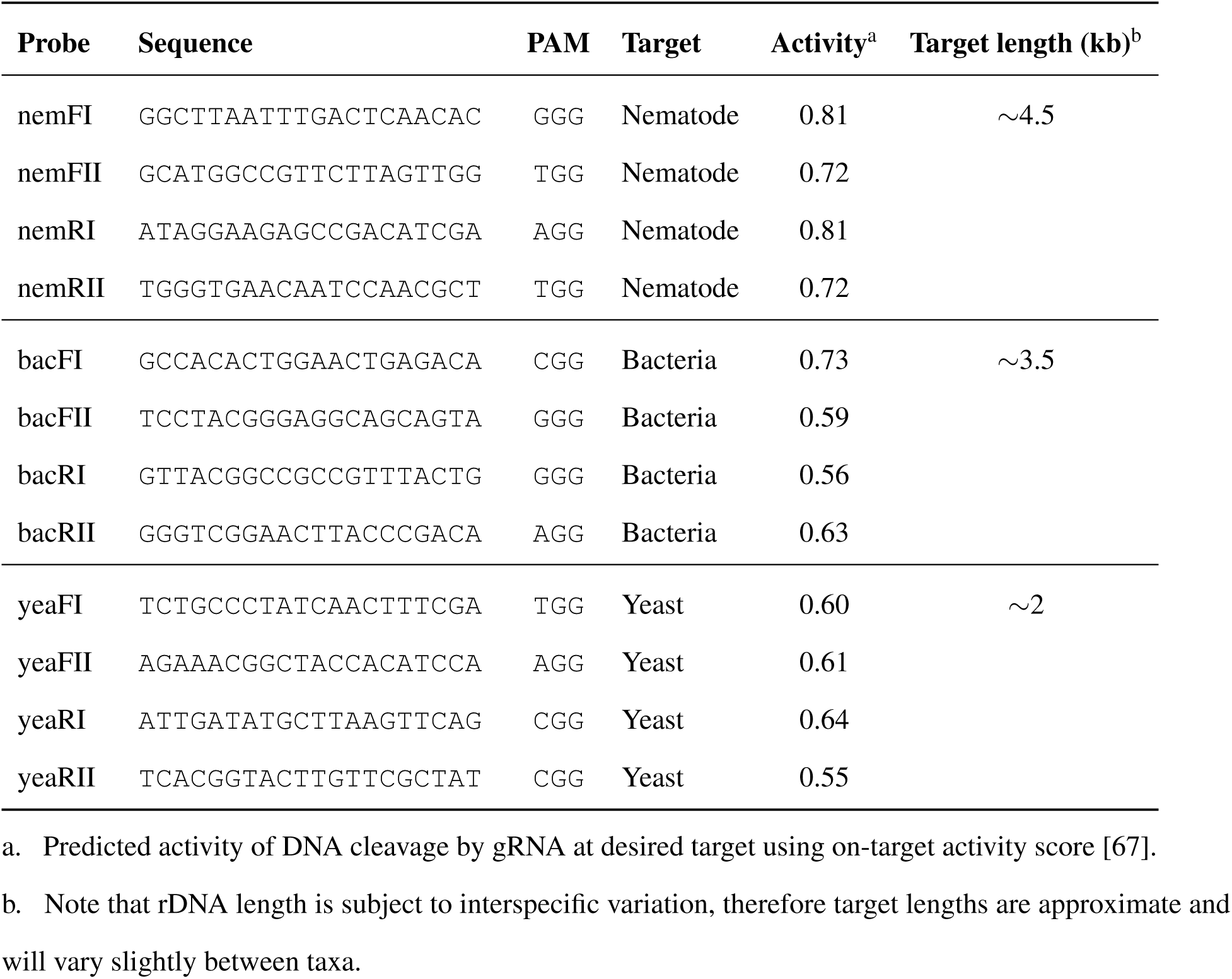
crRNA probes used for Cas9 cleavage of ribosomal DNA barcodes.

Analysis of the sequencing data using the metagenomics workflow estimated approximately 800 species in each sample, a gross overestimation of species richness compared to the ten species actually present in the mock community. However, most species from this report had an abundance of *<*1%. It was noted that nanopore R9.4.1 flow cells generate reads with an accuracy of 96.5% which is not sufficient to classify single reads to species level, and is the probable cause of the overestimated diversity. As a result, we decided to do the analysis to genus level. At the genus level, the metagenomics workflow estimated ∼380 genera, still a large overestimation, but all excess genera than those truly present had an abundance of ≤0.3%. The dominant ten genera in the metagenomics report matched the expected ten genera in close to expected proportions (**Data S3**).

To more precisely assess the ability of our enrichment method to capture accurate relative species abundance, we made use of the known rDNA copy numbers and genome sizes of each organism within the microbial community (**Table 1**). We carried out a quantitative analysis using an alignment workflow (**Methods**), providing the workflow with only reference sequences of the species present in the sample (the known ground truth), rather than the full NCBI reference database as in the metagenomics workflow. This helped us to remove confounding effects of the error-prone reads being compared to a very large reference database, inflating diversity estimates and possibly skewing relative abundance measures.

Agreement between the output proportions from our method and true community proportions (**Data S3**) was assessed using Lin’s concordance correlation coefficient (CCC) [52]. CCC assesses how well pairs of observations conform relative to another set, measuring both precision and accuracy. Observed versus expected proportions of each species exhibited high concordance, with Lin’s concordance correlation coefficient *ρ_c_* = 0.87 (*p* = 1.39 × 10^−10^, *t* = 9.67, *df* = 29) for combined data across three replicates (**Figure 5**). Yeast in our samples (*S. cerevisiae* and *C. neoformans*) were slightly over-represented in the output of all sequencing runs compared to bacteria, and *L. monocytogenes* was slightly under-represented (**Figure 5**).

**Figure 5:**
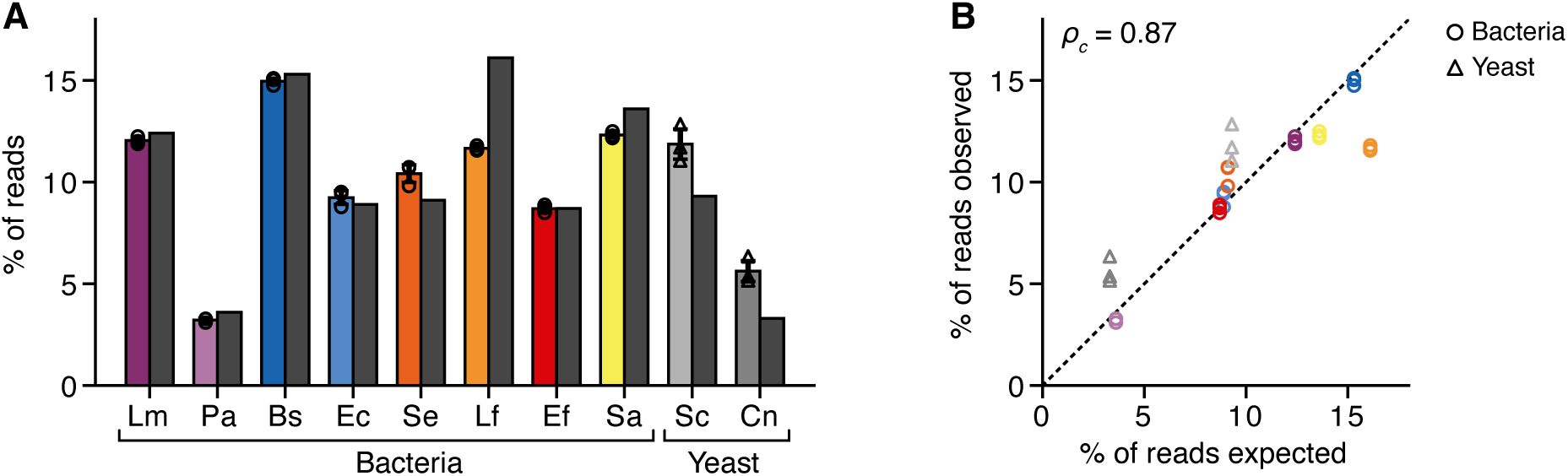
Analysis of a mixed microbial community using multiplexed crRNAs to target prokaryotic and eukaryotic species simultaneously. (**A**) Percentage of reads for each species in the ZymoBIOMICS Microbial Community DNA Standard (Lm: *Listeria monocytogenes*, Pa: *Pseudomonas aeruginosa*, Bs: *Bacillus subtilis*, Ec: *Escherichia coli*, Se: *Salmonella enterica*, Lf: *Lactobacillus fermentum*, Ef: *Enterococcus faecalis*, Sa: *Staphylococcus aureus*, Sc: *Saccharomyces cerevisiae*, and Cn: *Cryptococcus neoformans*). Coloured bars correspond to measured values from targeted sequencing. Individual replicate measurements shown by unfilled circles for bacteria and triangles for yeasts, and error bars show the standard deviation. Dark grey bars correspond to ground truth percentages of the species composition. (**B**) Comparison of observed and expected percentages of each species in the ZymoBIOMICS Microbial Community DNA Standard. Individual data points shown for each of the three biological replicates. Colouring of Data points matches bar colours in panel A. Dashed line denotes *x* = *y* and *ρ_c_* is Lin’s concordance correlation coefficient.

## DISCUSSION

In this study, we have demonstrated the ability to perform targeted nanopore sequencing using Cas9-based enrichment for the identification of nematodes and microbes in metagenomics samples without PCR amplification. High levels of enrichment of the target rDNA region in *C. elegans* demonstrates this is a feasible alternative to enrichment of barcodes using PCR amplification, which can present problems when attempting metagenomic analysis of samples. The method reliably identified nematodes, bacteria, and yeast to genus level with the false positives at very low abundance. A key feature of this method is the ability to multiplex crRNA probes to target diverse phylogenies in one reaction, an approach that is not possible with PCR primers due to the different temperature requirements of each primer during annealing. Moreover, the combination of the Cas9-based enrichment method with a long-read sequencing platform allows long DNA regions to be sequenced, which can increase taxonomic resolution by encompassing full-length or multiple barcode regions that capture greater phylogenetic variation [31, 32, 53]. In our study, not only did multiplexing allow qualitative analysis of a community, but also measures of relative abundance of a diverse metagenomic community consisting of both prokaryotes and eukaryotes.

Whilst our method was reliable at identifying up to genus level, there was a rich overestimation of taxa at species level. However this is a limitation of error-prone nanopore sequences rather than the Cas9-based enrichment approach. The average error rate of nanopore reads of 5–6% using R9.4.1 flow cells explains the generation of reads with false dissimilarity to species reference sequences. The newest R10.4 flow cells achieve read accuracies of over 99.1% [54], and so adaptation of our method for these flow cells would greatly increase the accuracy of taxonomic assignments to the species level. Higher accuracy reads from R10.4 flow cells would also mean more reads would pass quality filtering, allowing a fuller exploitation of the available data. Alternatively, with some modification to adapter ligation and computational pipelines, the Cas9-based enrichment approach to metabarcoding could be adapted to other long-read sequencing platforms, such as Pacific Biosciences (PacBio).

Whilst greater read accuracy would improve taxonomic assignment when target sequences have sufficient interspecific variation, it still holds that targeting short regions of a few hundred base pairs can limit delineation in closely related species [26]. We used an 18S reference database for taxonomic assignment of nematode sequences, but it has been shown that partial 18S sequences cannot distinguish between closely related *Caenorhabditis* species due to a lack of interspecific polymorphisms in that region [26]. Whilst even family level discrimination in nematodes can be sufficient for insight into ecosystem function [55], it is argued that species-level knowledge provides more information about community structure [56]. Furthermore, for other applications of metabarcoding, such as characterising human microbiome samples, delineation to species level is advantageous [57, 58]. Sequencing a longer region of multiple kilobases that spans full-length or multiple barcode regions can allow for greater taxonomic resolution [31, 33, 59]. Whilst we used an 18S reference database for nematode classification because of its compatibility with the EPI2ME bioinformatics workflow, our bioinformatics pipeline could be modified to maximally exploit our long-read data that spans small and large rDNA subunits, as well as ITS regions. Reference databases for each barcode of interest could be used together for cross-validation of reference sequences, as done by Heeger *et al*. for long-read barcoding of aquatic fungi using three databases simultaneously. This would provide better taxonomic resolution than using one single barcode [31].

The use of an amplification-free metabarcoding strategy bypasses difficulties associated with PCR amplification, such as problems amplifying low complexity regions, or stochastic variation early in the exponential process causing significant skewing of relative abundance estimates [21]. During PCR amplification, GC-rich regions are preferentially amplified due to differences in primer binding energies between AT and GC rich primers [21]. GC bias is strong in PCR primer-template hybridisation due to its sole reliance on binding between the primer and template, and is highly influenced by annealing temperature of the reaction. On the other hand, GC bias could be less pronounced in gRNA binding in CRISPR-Cas systems due to its additional PAM-site dependence and the stabilising effects of Cas proteins, and its lower sensitivity to temperature changes [60].

Similarly to PCR primer design, there were some challenges in finding highly conserved sequences targetable by crRNAs. There were some mismatches between crRNA sequence and templates. However, even relative abundance estimates did not seem significantly forfeited by the mismatches. This could be due to two reasons. First, the use of two gRNAs on each side of each target region might mean that if one sequence is not cleaved, the other gRNA complex acts in a redundant manner. This is supported by findings that using multiple gRNAs for each region of interest improve cleavage rates [46]. Alternatively, tolerance of mismatches by Cas9-gRNA complexes [61] might mean that the mismatches in our crRNA sequences and target genomes did not significantly affect binding and cleavage. Generally, PAM-distal mismatches have a small impact on cleavage of target DNA, whereas PAM-proximal mismatches tend to have a greater effect on binding and cleavage, the degree to which depends on the nucleotide substitution [61]. Baranova *et al.* [61] found that a C → G substitution on the first nucleotide upstream from the PAM affected cleavage time rather than the degree of cleavage, whereas a T → C and T → G substitutions reduced the rate and time of cleavage. So, whilst gRNA binding might not be subject to GC-bias when the probe sequence matches the template, when mismatches are involved, cleavage rates can vary. Our measures of relative abundance strongly matched true community proportions, but *L. fermentum* was one species that was slightly underrepresented. This might have to do with the presence of two mismatches between crRNA sequence and template in this species, in the middle of the crRNA, rather than one mismatch in other sequences or two PAM-distal mismatches in one crRNA-template combination. Increasing numbers of mismatches has been found to decrease the rate of cleavage [61]. Challenges around mismatches could potentially be addressed using crRNA sequences with degeneracy at a nucleotide position, as is sometimes done for PCR primers [62], or potentially extending the length of the cleavage reaction (**Figure 1C**).

Whilst some tolerance to template mismatches is a positive thing to allow crRNA sequences to bind to a greater diversity of sequences at one locus, this simultaneously means that off-target activity is also more likely. We encountered this problem when a sequence around 1.4 kb upstream of the reverse bacterial crRNA positions was likely cleaved by a fungal crRNA probe that has two mismatches and a suitable PAM site, shortening the expected ∼3.5 kb target region to ∼1.4 kb. Nevertheless, as full length reference sequences were provided to the analysis workflow, the remaining 1.4 kb of bacterial sequences after off-target cleavage left ample sequence to successfully map the shortened read against the reference and did not seem to impact qualitative or quantitative analyses. However, if the full target region is required for the delineation of closely related species or strains, one should be aware of possible off-target sites within the region of interest when designing crRNA probes. Enzymatic specificity is often dependent on enzyme concentrations, amongst other reaction conditions, so future work could be done to optimize enzyme/gRNA complex concentrations depending on the amount of starting DNA material to minimize off-target activity. Off-target activity outside the region of interest should not cause an issue, even if they are in a suitable orientation to cause extra reads to be generated, as sequences from other parts of the genome would be sufficiently divergent so as not to be mistakenly mapped to reference marker barcode sequences.

We found that our results were robust to varying amounts of starting DNA, ranging from 510–2917 ng in metagenomic samples. The number of reads generated from sequencing runs increased with increasing mass of starting DNA, but observed proportions of each species remained consistent across replicates. DNA yield that can be obtained by extraction from crude environmental samples can vary greatly [63], so testing the robustness of this method to smaller amounts of starting material would be beneficial for further method validation. Also, it would be interesting to test the detection limit of Cas9-enrichment, by executing similar experiments on mock communities with a larger range of abundances, e.g., using the ZymoBIOMICS microbial community standard with log distribution that contains species in abundances varying from 0.000089% to 89.1%.

In the current study, sequencing was run for up to 72 hours to ensure maximal data was generated. However, read generation was typically exhausted after 12-24 hours, indicating that such long sequencing times are not necessary. Knot et al [37] sequenced nematode DNA on a MinION for only ten minutes with accurate species assignment, whilst Hall and Speed [59] suggest 1 Gb of sequence data is appropriate for ultra-long read bacterial species identification. The optimal trade-off between sequencing depth and cost will depend on the complexity of the community being sampled and the length of the target region. If the community of interest requires a short sequencing time, combining this with the fast library preparation of Cas9-based enrichment (under two hours) could be used for rapid diversity and relative abundance assessments, bypassing the time-consuming PCR amplification process used in other metabarcoding approaches.

In summary, we show that it is possible to apply Cas9-based enrichment for taxonomic classification and relative abundance measures of metagenomic samples. The ability to multiplex crRNAs to target diverse phylogenies, combined with long-read sequencing technology to increase taxonomic resolution, gives this method great potential for characterising highly diverse biotic communities. It could also be used for rapid diversity assessments, bypassing the need for time-consuming PCR amplification. Adaptation of our approach to updated flow cell chemistry and powerful computational pipelines will enhance its species-delineation power, ensuring more in-depth assessments of organismal diversity.

## METHODS

### Organisms used in the study

Strains of *Caenorhabditis elegans* (AA1), *Caenorhabditis remanei* (JU724), *C. drosophilae* (DF5112) and *Panagrellus redivivus* (MT8872) were obtained from the Caenorhabditis Genetics Centre (CGC). *Heterorhabditis bacteriophora* and *Steinernema feltiae*, entomopathogenic species commonly used as a biological control agents in horticulture, were bought in the form of Nemasys Biological Chafer Grub Killer (BASF) and Nemasys No Ants (BASF), respectively, containing live infective juveniles. The ZymoBIOMICS Microbial Community DNA Standard (Zymo Research, D6306) was used for testing the method on a diverse set of microbes.

### Growth conditions

Nematode strains from the CGC were cultured on nematode growth medium (NGM) plates with OP50 *Escherichia coli* for eleven days at 25°C using standard methods [64].

### crRNA design

The use of multiple guides on each side of the region of interest has been demonstrated to improve cleavage rates [46], therefore, two crRNAs were designed for each side of the region of interest (a total of four probes for each target region). In total, twelve crRNAs were designed; four targeting nematode rDNA, four targeting bacterial rDNA and four targeting yeast rDNA (**Table 2**). crRNAs were ordered from Integrated DNA Technologies (IDT).

To design the nematode crRNAs, the position of the rDNA cluster in *C. elegans* genome WBcel235 (RefSeq GCF 000002985.6) [65] was located using the UCSC genome browser [66]. Within this region, the CRISPR target track on UCSC genome browser was used to search for DNA sequences with NGG PAM sites targetable by the Cas9-gRNA complex [66]. Target sequences were selected based on a combination of an efficiency score [67], specificity (uniqueness of 20-mer sequence in the genome) [68], and the conservation of bases across species as displayed on the Multiz Alignments and Conservation track on the UCSC genome browser [66]. Highly conserved sequences were selected to maximise phylum-wide annealing **(Figure 2)**. The region of interest between the custom crRNA guides was ∼4500 bp long, capturing partial 18S (506 bp) and 28S (2980 bp) rRNA genes, full 5.8S rRNA gene and full ITS1 and ITS2 regions (**Figure 2**). The position of our nemFII crRNA probe overlaps with the position of ‘NF1’ forward PCR primer site used for 18S nematode barcoding [69]. crRNA design was based on the *C. elegans* genome, and for other species present in the community that do not have annotated genomes, 18S and 28S sequences were aligned to determine the number and position of mismatches between crRNA and template for each species (**Figure 2**; see **Data S1** for sequence accession numbers).

The microbial mock community consisted of both bacteria and yeast species. To test the possibility of multiplexing crRNA probes for multiple targets in a single reaction, we designed two sets of crRNA probes to target the two separate taxonomic groups (**Table 2**; **Figure 2**). We designed bacterial crRNA probes that capture ∼1430 bp of 16S rDNA region in bacterial genomes (**Figure 2**), 16S being the most commonly used sequence for bacterial barcoding [70]. We also designed crRNA probes to target the yeast species present in the mock community, capturing the full nuclear ITS region **(Figure 2)**, widely used in fungal barcoding [71]. The final target regions also included other barcode regions that can be used for taxonomic classification, such as the ITS and partial 23S region for prokaryotes, and partial 18S, partial 28S, and ITS2 regions in eukaryotes (**Figure 2)**.

Genomes for each species (**Data S1**) were loaded into Geneious Prime version 2025.0 and the 16S or ITS regions were aligned for the bacteria and yeast, respectively. The ‘Find CRISPR sites’ tool was then used in Geneious, and CRISPR sites with the best combination of cutting efficiency scores and conserved bases across the species were chosen, whilst maximising the length of the target barcode included within the cleavage sites (**Table 2**; **Figure 2**).

### DNA extraction

DNA extraction from nematodes was performed using the Monarch Genomic DNA purification kit (New England Biolabs, T3010S) and the tissue extraction protocol. Samples underwent ethanol precipitation to concentrate DNA. NaOC was added in 0.1 volumes to each sample, followed by 2.5 volumes of absolute ethanol, and incubated at –20°C for 30 min. Tubes were centrifuged at 14,000 × *g* for 30 minutes at room temperature and supernatant was removed and discarded. Samples were rinsed with 70% ethanol and centrifuged at 14,000 × *g* for 15 min. The supernatant was removed, the pellet air dried and then resuspended in 0.2 volumes of nuclease-free water. DNA quality was checked on a NanoPhotometer NP80 spectrophotometer (Implen). DNA in each sample was quantified using the Qubit broad-range dsDNA assay kit on a Qubit 3.0 fluorometer (Invitrogen). The molecular weight of the DNA was checked on a 1% agarose gel stained with GelGreen (Biotium, BT41005), loaded with 6X purple loading dye (NEB), using the 1 kb plus DNA ladder (New England Biolabs, N3200S). High molecular weight samples were retained for subsequent steps. DNA for the microbial mock community was purchased as a ready-to-use DNA mixture (Zymo Research, ZymoBIOMICS Microbial Community DNA Standard, D6306).

### Library preparation and nanopore sequencing

Library preparation was done using the Cas9-based enrichment ligation sequencing kit (Oxford Nanopore Technologies, SQK-CS9109) following manufacturer’s protocol. In total, twelve libraries were prepared and sequenced independently (**Data S2**). Libraries were loaded onto a R9.4.1 flow cells (Oxford Nanopore Technologies, FLOMIN106D) and sequencing performed on a MinION Mk1B device using MinKNOW version 22.12.7.

To investigate the performance of the crRNA probes and the level of enrichment that can be achieved using this approach, the method was applied to a single nematode species, *C. elegans*. A single species was used in initial tests due to ease of mapping sequencing reads to a single genome, simplifying the analysis of enrichment efficiency and target region coverage. Nematode crRNA probes (**Table 2**; **Figure 2**) were used in library preparation. Three separate libraries were prepared (Samples 1a–c) with input gDNA quantities between 1400 ng and 2500 ng (**Data S2**), and these were sequenced independently for ∼20 hours.

Next, the method was applied to DNA samples containing a mixture of three nematode species (Samples 2a–c; **Data S2**); *C. elegans*, *H. bacteriophora*, and *S. feltiae*, to assess whether this method can identify multiple species in metagenomic samples. Again, nematode crRNA probes (**Table 2**) were used in library preparation. Using the same approach, this method was then applied to a mock community containing DNA from six nematode species (Samples 3a–c; **Data S2**); *C. elegans*, *H. bacteriophora*, *S. feltiae*, *C. remanei*, *C. drosophila* and *P. redivivus*. Total starting DNA mass ranged from 510–2917 ng for each mock nematode community sample (**Data S2**).

Finally, to assess whether this method could be used to quantify diverse taxonomic groups in a single library preparation step, eight crRNA probes targeting both bacteria and yeast genomes (**Table 2**) were applied to a microbial mock community containing DNA from ten microbial species (ZymoBIOMICS Community DNA Standard, D6306). These eight crRNA probes were added in equimolar concentrations during library preparation as per the manufacturer’s standard protocol.

### General bioinformatics

Base calling was performed using Guppy version 6.3.8. Base called reads in the FASTQ format were then filtered in Geneious Prime version 2025.0 such that Q *>* 10. Analysis of base called reads was done using EPI2ME version 4.1.3 functioning with Docker desktop version 4.17.0.

### Cas9-enrichment EPI2ME workflow

To analyse enrichment performance on a single genome, nanopore reads from the single species experiments (Samples 1a–c) were analysed using the wf cas9 workflow version 0.1.9 [72]. The full genomic sequence from *C. elegans* WBcel235 was used as the reference sequence (RefSeq GCF 000002985.6) [65, 73] to map nanopore reads to. The ‘target region’, used for analysing on-target versus off-target reads, was defined as the region between the two innermost crRNA probes on either side of the region of interest (genomic coordinates chrI:15063241–15067815).

### Metagenomics EPI2ME workflow

To assess the ability of our method to detect multiple species from metagenomic samples, the EPI2ME wf metagenomics v2.11.0 workflow was used [72]. The workflow was run using the minimap2 sub-workflow to allow for taxonomic classification of reads by mapping reads against a reference database. On the EPI2ME Labs worfklows, the reference can be chosen from a selection of databases covering archaeal, bacterial, and fungal data. Minimum match between read and reference was set to 98%.

For our analysis of the microbial data, the NCBI targeted loci database including 16S, 18S, 28S, and ITS sequences from archaea, bacteria, and fungi, was selected. For analysis of metagenomic nematode data, custom reference files were created because the default EPI2ME databases do not include eukaryotic sequences, aside from fungi. We created a custom sequence database and mapping file for use with the wf metagenomics workflow. This made use of the reference list from 18SNemaBase (18S NemaBase.fasta) [74] and generated the ref2taxid 18S NemaBase.txt file that maps the species of each reference to a valid taxid in NCBI. This is possible by using the taxid2name.txt file from taxonomy exports from the NCBI. Any references not present in the NCBI mapping were given the taxid 28384 (“other sequence”). For this database to be used in the wf metagenomics workflow, it was necessary to select the minimap2 aligner, using the 18S NemaBase.fasta file as the reference library and ref2taxid 18S NemaBase.txt as the ref2taxid mapping option. Scripts used to generate the database and mapping are available at: https://github.com/BiocomputeLab/18SNemaBase-EPI2ME.

### Alignment EPI2ME workflow

The EPI2ME wf alignment workflow version 0.3.3 [72] was used for quantitative analysis of microbial metagenomic sequences. Full length target sequences of each species in the mock community were extracted from genomic sequences (**Data S1**) and provided to the workflow as references to map nanopore reads against. Coverage of each reference sequence after mapping was used as the relative abundance data.

### Statistical analysis

A Lin’s concordance correlation coefficient (*ρ_c_*) was performed using RStudio version 4.4.1 to assess how well the observed proportions compare to the true proportions of each species in the microbial mock community. Data from three replicates were combined into one dataset, and a single coefficient was calculated for the combined quantitative data.

## Supporting information

Data S1 - Species Accession Numbers

Data S2 - Sample Compositions and Sequencing Details

Data S3 - Quantitative Results for wf_alignment and wf_metagenomics

## DATA AVAILABILITY

**Data S1** contains the accession number for genomic sequences used for crRNA design and as reference sequences for mapping reads against. **Data S2** provides information on the content and composition of each library that was used for sequencing. **Data S3** outlines results for quantitative analysis workflows. All basecalled nanopore sequencing data and a snapshot of the custom 18S NemaBase reference database for EPI2ME is available at Zenodo: https://doi.org/10.5281/zenodo.14250758

## ACKNOWLEDGEMENTS

This work was funded by a Liv Sidse Jansen Memorial Foundation grant (to T.E.G., C.C. and R.C.) In addition, T.E.G. was supported by a Royal Society University Research Fellowship grant URF\R\221008, a Turing Fellowship from The Alan Turing Institute under EPSRC grant EP/N510129/1, and the UKRI Engineering Biology Mission Award CYBER under BBSRC grant BB/Y007638/1. The funders had no role in study design, data collection and analysis, decision to publish or preparation of the manuscript. Some nematode strains were provided by the Caenorhabditis Genetics Centre, which is funded by the NIH Office of Research Infrastructure Programs (P40 OD010440).

## AUTHOR CONTRIBUTIONS

T.E.G., C.C., R.C. and L.N.-R. conceived the project. L.N.-R. carried out all experiments, developed the methodology, and wrote the initial draft of the manuscript. C.C. and R.C. assisted with experiments. All authors contributed to the data analysis, troubleshooting of experimental methods, and editing of the manuscript.

